# Single-cell RNA Sequencing Analysis of Sputum Cell Transcriptomes Reveals Pathways and Communication Networks That Contribute to the Pathogenesis of Asthma

**DOI:** 10.1101/2025.03.31.646405

**Authors:** Xiting Yan, Qing Liu, Taylor Adams, Jonas Schupp, Sihan Li, Stark Huang, Nicole Grant, Gabriella Wilson, Jose Gomez, Lauren Cohn, Naftali Kaminski, Scott T. Weiss, Kelan Tantisira, Geoffrey L Chupp

## Abstract

**Background:** Asthma is driven by complex interactions amongst structural airway cells, cells of the immune system, and the environmental. While sputum cell characterization has been instrumental in studying asthma pathogenesis and refining treatment strategies, the nuances of cellular transcriptomes and intercellular communication in asthmatic sputum remain poorly understood.

**Methods:** We employed single-cell RNA sequencing to analyze cells isolated form the sputum from 16 asthma patients and 8 non-asthmatic controls. Cell identities were established using curated marker genes and SingleR annotation. We compared cell-specific gene expression and communication networks between asthmatic and control groups, correlating findings with distinct pathways that were dysregulated in asthma.

**Findings:** 37,565 cellular transcriptomes were captured and analyzed. 15 distinct cell populations were identified, including various macrophages, monocytes, dendritic cells, and lymphocytes, along with rare cell types such as mast cells, innate lymphoid cells, bronchial epithelial cells, and eosinophils. Intercellular communication analysis indicated heightened signaling activity in asthma compared to controls, particularly in CD4+ T cells and dendritic cells which exhibited the most significant increases in RNA expression of outgoing signaling molecules. Notably, the ADAM12-SDC4 and CCL22-CCR4 ligand-receptor pathways demonstrated the strongest shifts between asthma and control subjects, particularly between dendritic cells and CD4 lymphocytes.

**Interpretation:** SC RNA seq profiling the asthma cellular transcriptome analysis of sputum highlights both innate and adaptive immune mechanisms that are significantly amplified in asthma. The elevated expression of ADAM12-SCD4 and CCL22-CC4 point to their critical role in asthma pathogenesis, suggesting potential avenues for targeted therapies and improved management of this chronic condition.

**Research in context:** *Evidence Before This Study:* Asthma is a chronic inflammatory disease of the airways driven by intricate interactions between airway structural and immune cells. Previous transcriptomic studies have focused on bulk RNA samples from the airway, leaving significant gaps in our understanding of the cellular dynamics that characterize the disease.

*Added Value of This Study:* This study pioneers the use of single-cell RNA sequencing on sputum samples from patients with asthma, revealing a detailed landscape of cell phenotypes and dynamic communication patterns that distinguish asthmatic individuals from those without the disease. Notably, heightened intercellular communication was observed in asthma, particularly between CD4+ T cells and dendritic cells, confirming that there is a robust network of interactions between immune and structural cells. The notable increase of ADAM12-CCR4 communication from dendritic cells to other cell populations further emphasizes the dysregulation present in asthma.

*Implications of All Available Evidence:* Our transcriptomic profiling illuminates distinct and amplified communication pathways involving CD4+ T cells and dendritic cells, aligning with established paradigms of both adaptive and innate immune responses in asthma pathogenesis. The identification of ADAM12 and CCR4 pathway dysregulation adds a critical layer to our understanding of the molecular mechanisms underpinning asthma, paving the way for potential therapeutic targets and personalized treatment strategies. Single cell profiling of the sputum has the capacity to characterize the breadth of cellular phenotypes, their functional status, and the communication in the airway at a level not previously attainable.

## INTRODUCTION

Our current understanding of asthma pathogenesis proposes that chronic inflammation of the airways results from dysregulated interactions between structural cells of the airways, cells of the immune system. There is now clear evidence that interactions between the structural cells of the airway and resident immune cells, in particular, dendritic cells and T cells through receptors and secreted mediators impact the heterogeneous clinical expression of the disease. Targeted inhibition of these mediators can dramatically improve disease control and may have the potential to be disease modifying if implemented at the appropriate time in the course of disease and in the right subtype of patient. Achieving this lofty goal will require precise characterization of the cellular interactions in patients with asthma and clinical studies of how targeted therapies affect these interactions. Reliance on non-human models of the disease in species in which asthma does not occur naturally and analysis of mediator expression in samples isolated from tissue biopsies, brushes, bronchoalveolar lavage fluid, or cell pellets has to date made this difficult. The latter techniques are limited by their invasiveness and bulk nature that often averages signals across samples and study size that limits the generalizability of the results. In addition, cell-based assays by immunohistochemistry, in situ hybridization, or FACS provide cell-based expression. But the number of mediators is limited so, multidimensional computational analyses are not feasible.

In contrast to these approaches, single cell RNA sequencing (scRNA-seq) analysis of cells isolated from the airway provides a revolutionary opportunity for asthma research. First, it provides high dimensional data from each cell that can be used to define a wide range of cell types as well as their functional status. Second, scRNA-seq eliminates the averaging of transcriptomic signals created by analysis of proteins and mRNA isolated from a cell pellet, so called “bulk” analysis. Third, because it is cell-based, this method can be used on samples with small numbers of cells, a feature particularly valuable for analysis of sputum samples where yield from patient to patient can vary significantly^1^. Fourth, high dimensional cellular transcriptomes can be used to infer communication amongst cell types using computational techniques which is not possible using bulk RNAseq methods. Thus, scRNA-seq of cells isolated from the airway non-invasively represents a major opportunity for the cellular immunologic study of asthma and other chronic airway diseases.

To date, several groups have applied scRNA-seq to study lung disease, and in particular asthma. These studies included analysis of samples from bronchoalveolar lavage fluid (BAL)^2^, endobronchial brushing^3^, airway biopsy ^4,5^, lung tissue^6,7^, peripheral blood mononuclear cells (PBMCs)^8,9^, and airway epithelial cells^10,11^. The single-cell studies of BAL^2^, endobronchial brushing^3^, lung tissue^6^ and PBMC^8,9^ profiled cells from flaring or stimulated samples instead of steady-state samples, lacking the ability to identify changes in patients with asthma at baseline. The lung tissue single-cell study by Everman et al.^7^ profiled cells from healthy donor lungs and thus do not reflect changes in patients with asthma. The airway epithelial cell single-cell studies^10,11^ profiled cultured differentiated human airway epithelial cells and airway epithelial brushing samples after allergen challenge. The endobronchial brushing single-cell study profiled cells from allergic asthmatics and allergic non-asthmatic before and after segmental allergen challenge (SAC) with a focus on identifying responses to SAC in allergic asthmatics (AA) compared to allergic controls (ACs). The airway biopsy single-cell study profiled steady-state samples and captured the most comprehensive panel of cell types in the human airways, including epithelial, stromal and immune cells. Compared to these previous single-cell studies in asthma, our study conducted single-cell analysis of induced sputum samples which was extracted from human airways under the steady-state using a non-invasive way. Our data captured similar panel of cell types to the previous studies except for the stromal cells in the airway biopsy samples. Part of our findings are also overlapped with those in previous studies.

In this study, as a follow-up to our previous publications on scRNA-seq^1,12^, we demonstrate the feasibility of scRNA-seq on cells isolated from the sputum from non-asthmatic controls and patients with asthma and optimize it for characterization of sputum eosinophil transcriptomes. Using a protocol optimized for preservation of RNA, we identified at least 15 different cell populations including multiple subtypes of immune, resident, and structural cells of the lung. We also demonstrate how gene signatures differ and how the cell-to-cell communication networks differ between asthma and controls and some of the pathways mostly strongly associated with these differences. We further optimized the protocol and identified eosinophils from both blood and sputum samples.

## METHODS

### Yale Center for Asthma and Airway Diseases (YCAAD) Cohort

Sputum samples were collected from asthmatic and control subjects that completed the Yale Center for Asthma and Airway Diseases (YCAAD) phenotyping protocol previously described.^13^ Briefly, inclusion criteria for enrolled subjects were: ≥ 18 years of age, non-smoking, and a history of asthma and pulmonary function testing consistent with a diagnosis of asthma based on published guidelines.^14^ Exclusion criteria included smoking within the past year, a history of chronic lung disease other than asthma (i.e. chronic obstructive pulmonary disease, pulmonary vascular disease or interstitial lung disease); other severe chronic conditions that would limit the subject’s ability to safely undergo the required procedures. Subjects were administered an asthma history questionnaire that included an asthma control test (ACT) score. Spirometry was conducted before and after short-acting bronchodilator administration and exhaled nitric oxide (FeNO) was measured in adherence with the American Thoracic Society guidelines.^15^ Sputum induction was performed with hypertonic saline, as previously described.^16–19^ Mucus plugs were dissected from the sputum sample and the cellular and aqueous compartments were separated using DTT. Total cell count, cell viability and differential, were determined by hemocytometer, trypan blue exclusion, and Wright-Geimsa stain, respectively, and the cell pellet was processed or stored in DMSO as previously described.^20^ For single cell RNA sequencing analysis, the sputum cell pellets were re-suspended and processed using the Chromium V2 chips or the Flex Fixation kit per the 10x Genomics protocol (Boston, MA). The white blood cells were suspended and processed using the Flex Fixation kit. The resulting single cell cDNA library was sequenced on an Illumina HiSeq2000. The data can be obtained from the GEO database (GSE270863).

### Computational analyses

Raw sequencing reads were preprocessed using the Cell Ranger Pipeline V3.0.1 for demultiplexing, cell barcodes error correction, background GEM removal and data quality assessment. Reads from high-quality samples were further mapped to the human genome with annotations for both mRNAs and non-coding RNAs using STAR^21^. The number of unique molecular identifiers (nUMI) was calculated for each gene in each cell based on mapping results, which were further processed using the Seurat^22^ R package (V3.0) for data visualization, cell clustering and marker gene identification. Cell lineage of each cell cluster was identified and assigned using manual classification^5,12,23^ and SingleR^24^ R package (V2) with the human primary cell atlas (HPCA)^25^ as the reference dataset. Squamous epithelial cells were separated from airway epithelial cells based on canonical marker genes for squamous cell markers (Supplemental Materials) and excluded from downstream analysis. Differentially expressed genes between asthma and controls in each cell type were identified using the Student’s T test on the average gene expression level across cells from the same cell type per sample. Genes with a nominal p value < 0.05 and fold change > 2 were considered significant. Pathway enrichment analysis was conducted on the identified DEGs using MetaCore and pathways with false discovery rate (FDR) less than 0.05 were considered significant. Cell-cell communication networks were built in each subject separately using Connectome^26^ based on the expression levels of ligand-receptor (LR) pairs defined in the Fantom5 database. The outgoing and incoming centrality of each cell type were evaluated and compared using the Student’s t test between asthma patients and controls to identify differences in communication among cell types between asthma and control. The expression of each LR pair was also compared to identify LR pairs driving the communication changes within a given signaling family. For both comparisons, a nominal p value < 0.05 was considered as significant.

## RESULTS

The study flow for the processing of the sputum samples can be found in Figure 1A. Clinical characteristics of the study subjects including demographics, asthma history, pulmonary physiology, and sputum cell differentials determined by light microscopy are shown in Table 1. From the 24 samples, a total of 37,565 cells were captured with an average of 1,174 cells and 78,534,771 reads per sample. On average, 15,532 genes per sample and a median of 1,203 genes per cell were measured with an average number of unique molecular identifiers (nUMIs) per cell of 4,186. A detailed summary of the sequencing data characteristics can be found in Supplementary Table S1.

**Figure 1.**
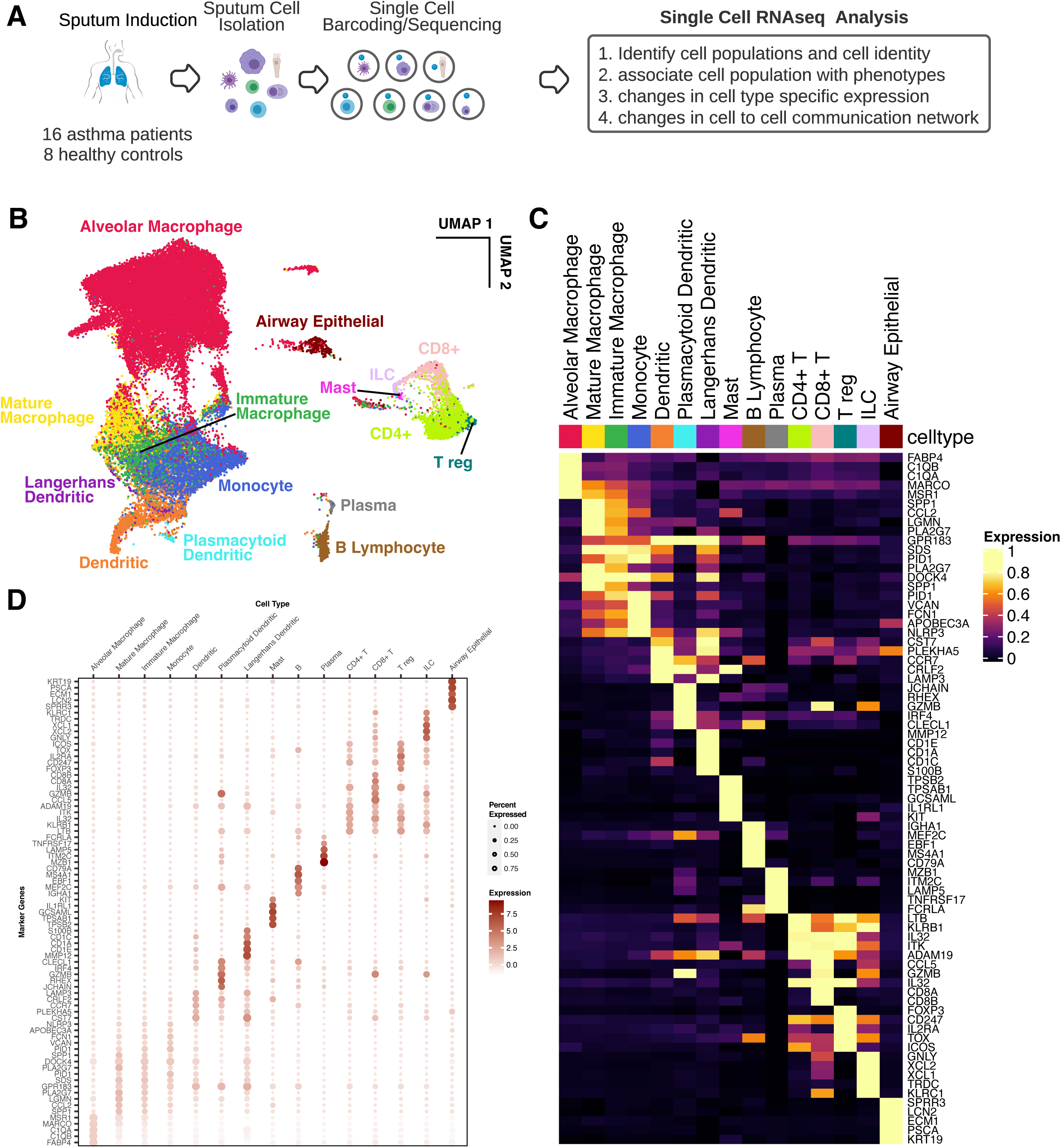
Cell populations identified in sputum. (**A**) Study workflow diagram. (**B**) Uniform Manifold Approximation and Projection (UMAP) of all cells classified into different cell types; each dot represents a single cell, and cells are labeled as one of 15 discrete cell types. (**C**) Heatmap of marker genes for the major myeloid and lymphoid cell lineages. (**D**) Heatmap of marker genes for cell types within the lymphoid cell lineages. (**E**) Heatmap of marker genes for cell types within the myeloid cell lineages.

**Table 1:**
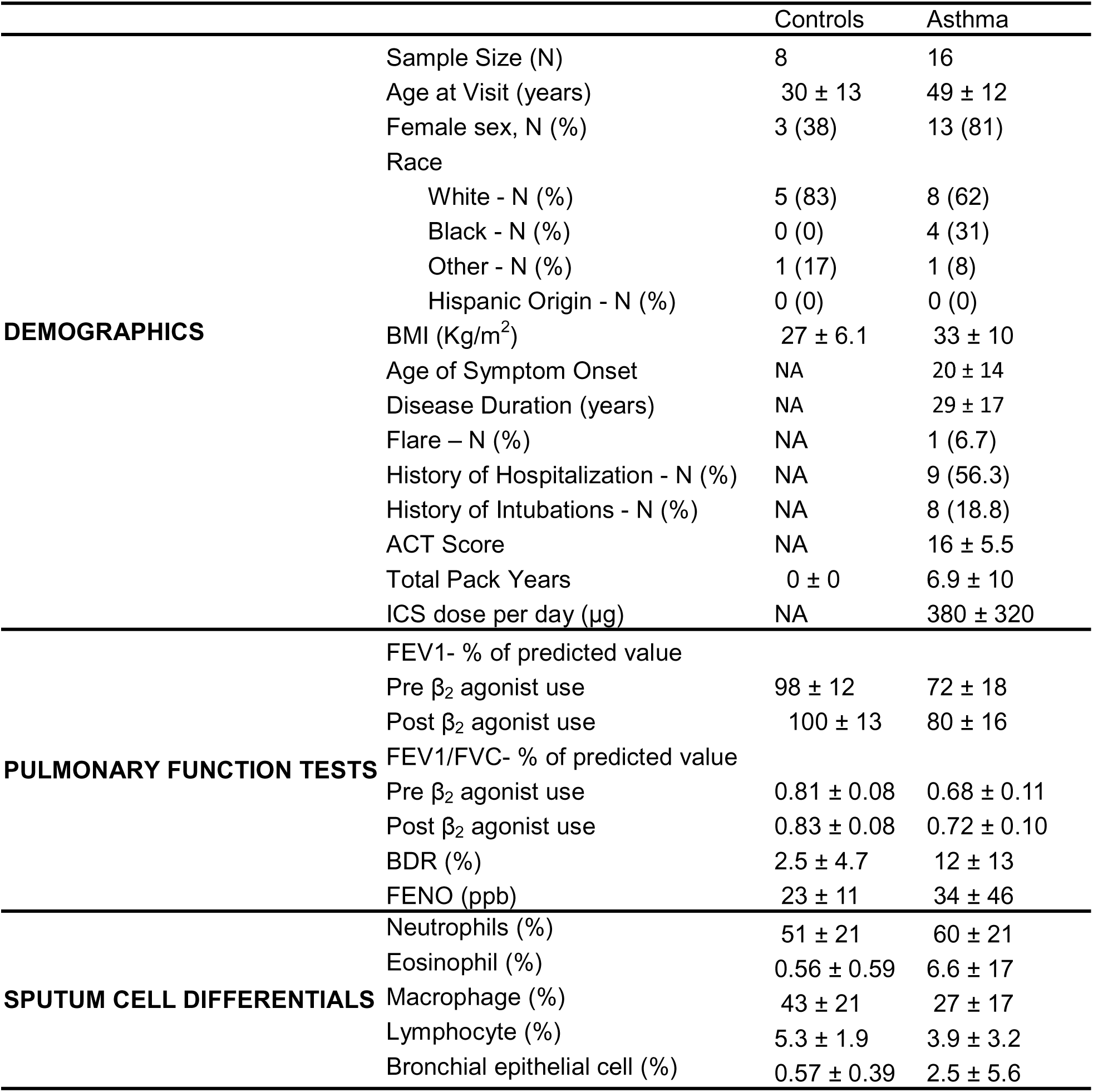
Demographics, pulmonary function tests and cell differentials of the 24.

### Cell populations identified in sputum

Integrated analysis of the data using Seurat identified 28 clusters of cells (Supplemental Figure S2B). By both manual classification using marker genes and correlation with the human primary cell atlas (HPCA) dataset there were 15 different known cell populations - mostly immune cells of myeloid and lymphoid lineage (Figure 1C). Marker genes of cell type (Figure 1D) matched those reported in other studies of immune cells isolated from the airways of patients with asthma and cystic fibrosis.^3,5,12^

Myeloid lineage cells included macrophages (71.8%), dendritic cells (5.1%), monocytes (12.7%), and mast cells (0.09%). Within the macrophage lineage, approximately 81.0% were immunophenotyped as alveolar macrophages based on increased expression of FABP4, C1QB, C1QA, MARCO, and MSR1 (Figure 1C-D). This subpopulation of macrophages showed the most variation in gene expression across individuals (Supplemental Figure S1). Of the remaining macrophage population (19% of the total), 43% of the cells demonstrated increased expression of genes associated with macrophage maturation and differentiation (SPP1, APOC1, ACP5, CTSS, GPNMB, APOE, CHIT1) indicating that they were mature macrophages while the other 57% were immature. Within the dendritic cell lineage, 4% of the cells were plasmacytoid dendritic cells based on increased expression of JCHAIN, RHEX, GZMB, IRF4, and CLECL1. 6% of the cells were Langerhans dendritic cells based on increased expression of MMP12, CD1A, CD1E, CD1C and S100B. The rest of the dendritic cells were the classical dendritic cells with higher expression of CCL22, CCR7, CRLF2 and LAMP3, and PLEKHA5. The analysis demonstrated the capacity of scRNA-seq to identify multiple types of myeloid cells in the sputum.

Lymphoid lineage cells were less common than myeloid cells and included T lymphocytes and Innate Lymphoid Cells (ILC) (7.7%), and B lymphocytes (1.6%). Within the T lymphocyte population, 67% were CD4+ helper T cells (CD40LG, SOS1, CCL20, and RBMS1), 28% were CD8+ cytotoxic T cells (CCL5, GZMB, CD8A, CD8B), 5% were T regulatory cells (FOXP3, IL2RA, CD247, TOX, ICOX), 5% ILCs (GNLY, XCL1, XCL2, TRDC, and KLRC1). Within the B lymphocyte population, 83% were B cells (IGHA1, MEF2C, EBF1, MS4A1, and CD79A) and the other 17% were plasma cells (MZB1, ITM2C, LAMP5, TNFRSF17 and FCRLA). Therefore, a broad array lymphoid cells were accessible via analysis of non-invasively collected sputum cells. (Supplemental Figure S3B).

There was a small percentage of airway epithelial cells (358 cells, 0.99%) and mast cells (31 cells, 0.09%). Granulocytes, including neutrophils and eosinophils, were not identified in freshly processed or DMSO preserved samples. This was likely due to low RNA levels and fragility of these cells, which limits GEM formation in the 10x flow cell and sequence tags in the 10x v2 kit. Four samples were also processed using the Flex Fixation kit from 10x Genomics, which enabled the capture of eosinophils and neutrophils. Sputum eosinophils were found to have increased expression of HDC, CCL, IL-5R (Supplemental Figure S7). Blood eosinophils expressed slightly different marker genes.

### Differential gene expression between asthma and controls

To understand how each cell type is functioning in asthma, differentially expressed genes (DEGs) were identified between asthma and controls and pathway enrichment analysis was conducted using the identified DEGs in MetaCore. Mast cells, plasma cells, and airway epithelial cells were excluded from this analysis because they were present in fewer than 3 subjects (Supplementary Table S2). Heatmaps of expression profiles of the identified DEGs for the 12 cell populations are shown in Figure 2. A detailed list of DEGs identified in each cell population can be found in Supplementary Table S3. Pathway enrichment analysis results of the DEGs can be found in Supplementary Table S4.

**Figure 2.**
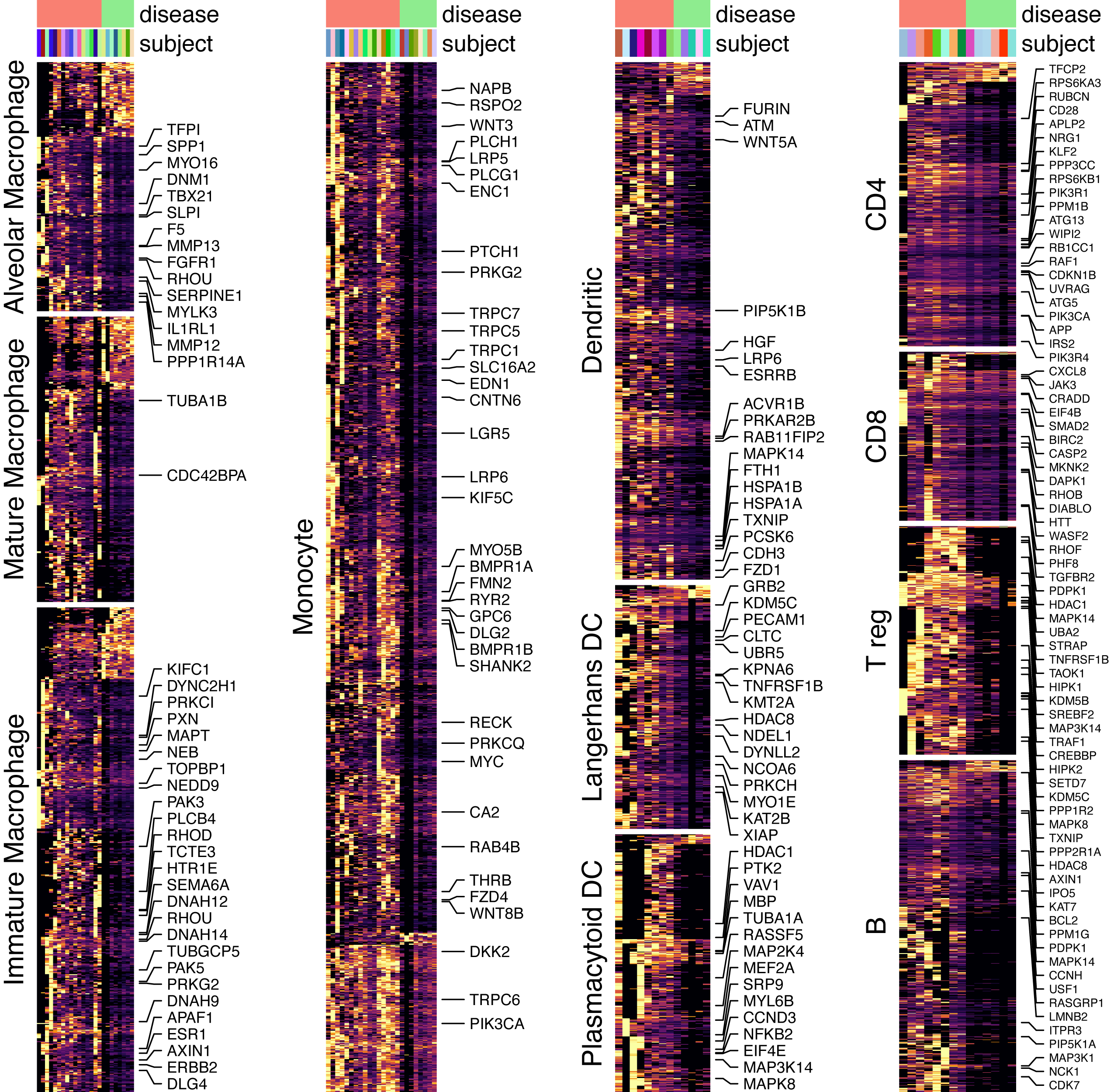
Differentially expressed genes between asthma patients and controls in each cell type. Heatmap showing expression profiles of the differentially expressed gene between asthma patients and controls in all cell types except mast cell, plasma cell and airway epithelial cells (significance P < 0.05).

Pathway enrichment analysis of DEG increased in alveolar macrophages revealed enrichment of tissue remodeling pathways and genes (SERPINE1, MMP12, MMP13, SLPI, GFFR1) and actin cytoskeleton remodeling (MYLK3, PPP1R14A, RHOU, and PRKAR18), indicating an active role of resident lung macrophages in structural changes associated with asthma^27,28^. Notably, IL1RL1, a receptor of the alarmin IL33 that promotes Th2 cell maturation^29^, and NLRP2 that inhibits NF-kB signaling^30^, were also increased in Alveolar macrophages in asthma. 431 DEGs were increased in immature macrophages and showed enrichment of actin cytoskeleton and microtubule formation pathways (Supplementary Table S4). Both pathways have been linked to airway remodeling previously^31^. There were 1,620 DEGs increased in classical monocytes, the largest number among all the cell populations examined. Pathway enrichment analysis of these DEGs demonstrated the strongest enrichment in the WNT/Beta-catenin pathway. This signaling pathway was also enriched in classical dendritic cells.

In Langerhans dendritic cells pathways associated with microtubule formation, cell migration, transcriptional activity pathways were enriched in DEG that were increased in asthma. Plasmacytoid dendritic cells, although rare, were enriched in immune response and cytoskeleton pathways. Combined, this analysis is consistent with the activation of macrophages and dendritic cells that promote inflammation in the airway wall.

DEG were also increased in CD4+ T and CD8+ T lymphocytes in asthma and showed enrichment of pathways associated with immune response activation, autophagy, cell proliferation, and apoptosis. B cells showed enrichment of the B cell receptor pathway, apoptosis and mitosis indicating increased activation and turnover in asthma. T reg cells were enriched for oxidative stress, the WNT/Beta-catenin pathway, apoptosis, and histone acetylation and methylation.

### Analysis of cell-cell interaction networks

To understand the cellular communication networks that were altered in asthma, we conducted cell to cell interaction network analysis using Connectome and the Fantom5 database.^26^ Connectome infers the strength of interactions between cells by examining the expression levels of ligand and receptor pairs between cells. The interaction strength of each cell population with all the other cell populations is summed and normalized as “centrality” to describe the importance of the given cell type in the communication networks. Both outgoing (expressing ligand or “sending”) and incoming (expressing receptor or “receiving”) centrality are calculated. Airway epithelial cells, mast cells and plasma cells were excluded from this analysis due to the small number identified. To identify asthma specific interaction networks and signaling families, outgoing and incoming centrality was compared between asthma and control samples (Figure 3A and Supplemental Figure S6). In general, intercellular communication was much more active in asthma compared to controls as the majority of significant incoming and outgoing centralities were higher in asthma. Specifically, outgoing centrality from CD4+ T cells, alveolar macrophages and dendritic cells was increased the most in patients with asthma (Figure 3A, Supplemental Figure S6) which is consistent with the activation of the adaptive immune response in asthma. Interestingly, incoming centrality was most significant in alveolar macrophages, mature macrophages, and CD4+ T cells highlighting the role of these cells as critical effector cells in adaptive and innate immune responses in asthma. The incoming and outgoing centrality that were lower in asthma compared to control subjects were minimal (Supplemental Figure S6).

**Figure 3.**
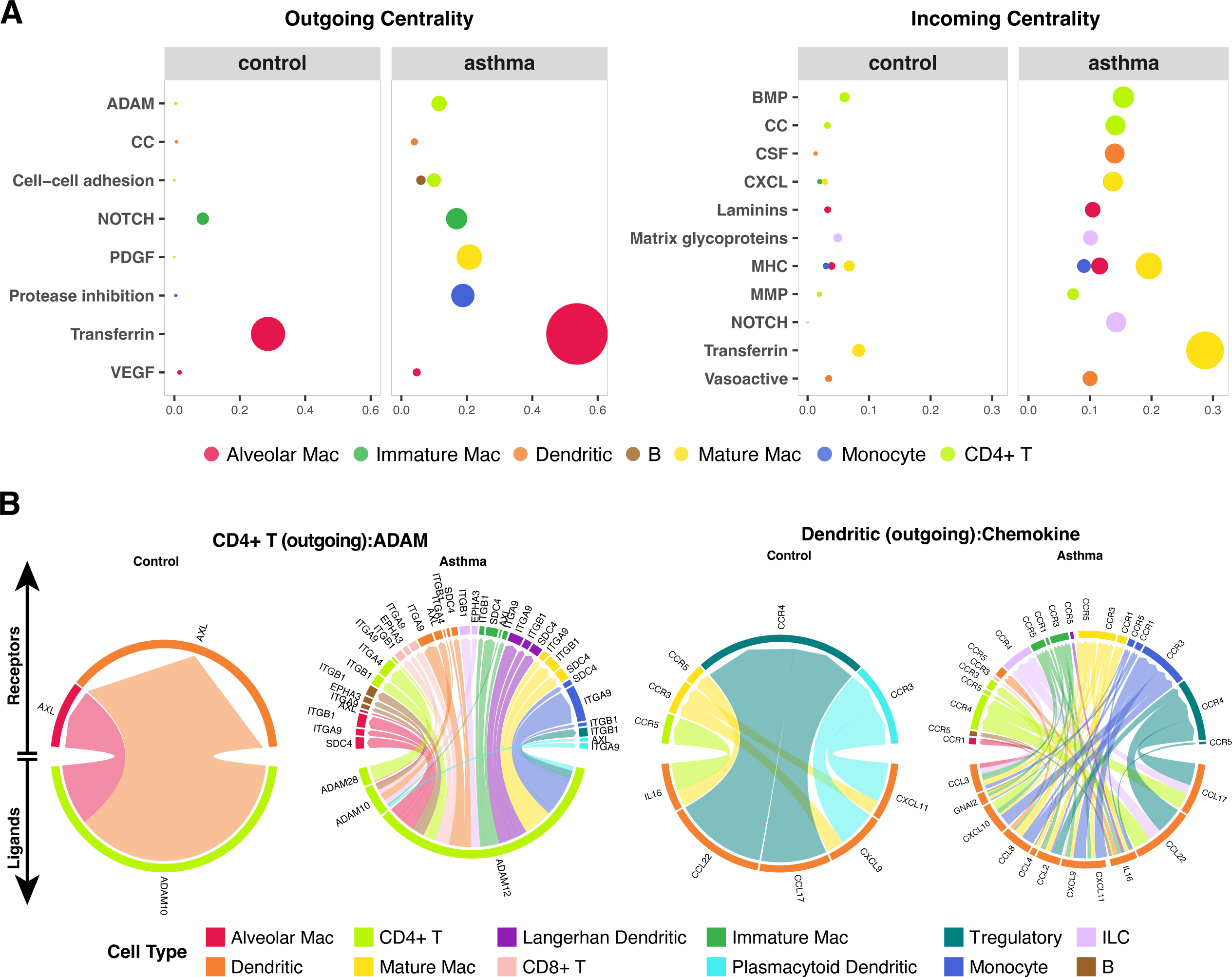
Differences of cell-cell interaction network between asthma and control samples. (**A**) Bubble plots demonstrating the change in outgoing centrality (left subpanel) and incoming centrality (right subpanel). In each subpanel, the outgoing or incoming centrality of different cell types (columns) across multiple signaling families (rows) was shown in control (left) and asthma (right). The size of the bubbles is proportional to the of the corresponding centrality value. For each cell type, only those signaling families with significant changes between control and asthma are shown. (**B**) Circus plot demonstrating the outgoing signals sent by CD4+ T cells for ADAM signaling family and dendritic cells for chemokine signaling family. The bottom half of the circle shows expression of the ligands and the top half shows the expression of receptors in different cell types (labelled by different colors). The width of the segments for each LR pair demonstrate its relative expression level in the corresponding cell type.

**Figure 4.**
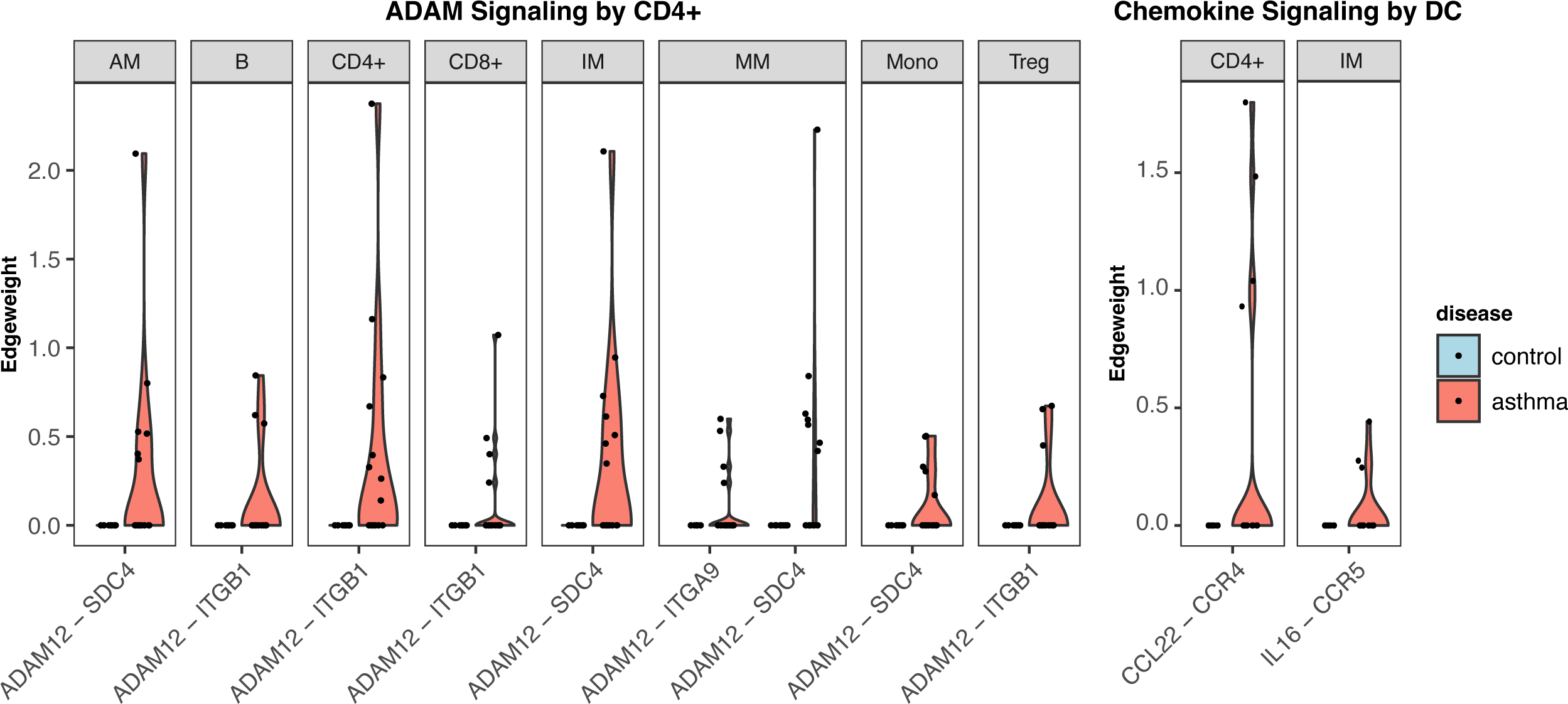
Ligand-receptor signaling differences between asthma and control. Violin plots of edgeweights for the significant ligand-receptor pairs from ADAM signaling family (sent by CD4+ T cells, left panel) and chemokine signaling family (sent by dendritic cells, right panel) were shown. Only LR pairs that were significantly (p<0.05) different are shown. Ligands expressed in CD4+ T cells (left panel) and dendritic cells (right panel). Receptors were expressed in cell types shown on the top of each violinplot. The edgeweight was calculated as the average log-transformed expression level in the ligand and receptor, representing the signaling activity level.

To identify the ligand and receptor pairs driving the higher activity in asthma, we focused on cell populations where the communication was most substantial-CD4+ T and dendritic cells. The most increased outgoing centrality from CD4+ T cells was from the ADAM receptor-ligand family (Figure 3B, Supplemental Figure S6) to macrophages and dendritic cells, and in an autocrine fashion to CD4+ T cells (Figure 3C). Specifically, ADAM10, ADAM 12, and ADAM28 were the ligands expressed in CD4+ T cells (Figure 3B). Comparison of the interaction strength of each ligand-receptor pair in the ADAM signaling family showed that ADAM12 and its receptors had the most significant difference between asthma patients and controls (Figure 3C). The receptors of ADAM12 included SDC4, ITGB1 and ITGA9, which were shown to be highly expressed in lung in the GTEx database^32^ matching the observations in this data. Moreover, expression of ADAM12 was significantly higher in asthma patients while expression of ADAM10 or ADAM28 in asthma and control was not significantly different between asthma and control. These results suggest that the ADAM signaling family, especially ADAM12, is dysregulated in CD4+ T cells in patients with asthma.

The professional antigen presenting cells and mediators of innate inflammation, dendritic cells had increased outgoing centrality of chemokines to immature macrophages, Tregs, monocytes and CD4+ T cells (Figure 3A-B), while the expression of CXCL10, CCL2, CCL4, CCL8, GNAI2, CCL3 or their receptors, was not present in the communication networks in control subjects (Figure 3B). CXCL10 from dendritic cells interacted with CCR3, CCR5 and CCR4 in CD4+ T cells and CCL4 from dendritic cells potentially interact with CCR1, CCR3 and CCR5 in mature macrophages. Specifically, the interaction between dendritic cells and CD4+ T cells through CCL22 and CCR4, and the interaction between dendritic cells and immature macrophages through IL-16 and CCR5 are both significantly higher in asthma compared to controls (Figure 3D).

Overall, there were more significantly increased incoming centralities than outgoing centralities (Supplemental Figure S6). Alveolar macrophage had increased laminins signaling suggesting that alveolar macrophages express extracellular matrix that contributes to airway remodeling. Other cell types with increased ECM expression included dendritic, Langerhans dendritic, immature macrophages, mature macrophages, CD4+ T cells, CD8+ T cells, Tregs, and B cells revealing a broad shift towards development of increased extracellular matrix that will sustain the influx of inflammatory cells in asthma. In comparison, monocytes, immature macrophages and mature macrophages were receiving signals from inflammatory chemokines, including CXCL and MHC molecules, consistent with their role in inflammation. Dendritic cells received higher colony-stimulating factors (CSF) signals, which regulates the survival, growth, differentiation and function of mononuclear phagocytes, granulocytes and myeloid progenitors, from CD4+ cells, alveolar macrophages, and mature macrophages. Innate lymphocyte cells (ILC) received higher NOTCH signals from immature macrophages, mature macrophages, and Tregs. Interestingly, CD4+ T cells received higher MMP signals from dendritic, immature macrophage, mature macrophage and monocyte. They also received higher Chemokine signals from alveolar macrophage, immature macrophage, mature macrophage, monocyte, dendritic and CD8+ T cells.

## DISCUSSION

The current paradigm of asthma pathogenesis suggests that alterations in the immune system and airway result in chronic inflammation, fluctuating airflow obstruction, and chronic symptoms that wax and wane. Efforts to determine the molecular basis of these alternations have been hampered by the heterogeneous nature of the disease, constraints of animal models, limited access to biopsies, and the poor resolution of traditional assays. Analysis of cells and supernatants isolated from mucus has proven to be a convenient, non-invasive, cost-effective method gain insights into airway inflammation. However, sample variability and yield have limited the scope of analyses. Here we introduce the single-cell RNA sequencing analysis of sputum samples from patients with asthma. We demonstrate the feasibility of this method on cells isolated from mucus, identify a broad array of cell types including granulocytes including eosinophils, and the capacity of this approach to reveal novel communication networks associated with asthma.

In contrast to other single cell transcriptomic studies of asthma to date, this is the first study of cells isolated from the sputum and adds to our previous study on single-cell RNA sequencing in Cystic Fibrosis. Our methodology shows the feasibility of single cell transcriptomic profiling with an input of 10,000 cells to yield thousands of cell transcriptomes. In addition, we also show that preserved samples and freshly processed samples have comparable transcriptomes (Supplemental Figure S2A), which has implications for the use of this methodology in multi-center studies where samples are banked and shipped for analysis in batches. In addition, the capture of on average 50,000 reads per cell allowed diverse phenotypes of cells to be identified including multiple subtypes of macrophages, dendritic cells, lymphocytes, and rare cell types such as ILC, mast cells, B-cells, bronchial epithelial cells, and granulocytes (neutrophils and eosinophils). Importantly, most of the cell populations were evident in similar percentages in both asthma and control patients, except for mast cells and bronchial epithelial cells which were rare, but predominantly present in the asthma patients. We also noted differences in gene expression in each cell type in asthma compared to controls with monocytes having the largest number of identified DEGs. Increased numbers of immune response genes were identified in asthmatic airways as expected and were pro-inflammatory and pro-airway remodeling in alveolar macrophages and immature macrophages.

The high dimensional nature of the single-cell RNA sequencing data allowed us to identify patterns in ligand and receptor expression that infers the activation of novel immunologic networks that are relevant to asthma pathogenesis. The CONNECTOME analysis revealed significantly higher centrality of multiple pathways in asthma compared to controls with outgoing centrality from resident immune dendritic cell populations to effector cells including T cells, B cells, and macrophages. Multiple receptor ligand pathways were significantly higher in asthma including the ADAM12-SDC4 signaling pathway and CCL22-CCR4 pathway. ADAM12 is a member of a metalloproteases family that mediates cell adhesion via a cysteine-rich domain syndecan-4 (SDC4) receptor interaction on the cell surface that triggers β1integrin-dependent cell spreading, stress fiber assembly, and focal adhesion formation^33^. The other ligand receptor pair includes the CC Motif Chemokine, Ligand 22 (CCL22), is expressed by dendritic cells and binds to CCR4 to attract Th2 cells in mice under bronchial challenge with house dust mite.^34^ The CCL22-CCR4 pathway was also shown to be increased in asthma in TH2 cells and dendritic cells in a single-cell RNA sequencing study of BAL samples from allergy challenged patients validating our findings and that these pathways are important mediators of both the acute allergic response and chronic inflammation associated with asthma^3^.

The promising results of this study also come with some limitations. First, the version of the 10x methodology we used was not able to capture the transcriptomes of eosinophils or neutrophils until it was modified and made available in the form of the 10x Prefix kit (Supplemental Figure S7). This highlights that granulocytes and in particular, eosinophils have low levels of mRNA, are extremely fragile and prone to lysis, have high levels of RNases. They have been extremely hard to identify by single-cell RNA-sequencing and have not been evident in other studies^5,6,35^. Second, the small of patients this study limited our ability to make correlations between rare cell types and asthma phenotypes, such as bronchial epithelial cells, mast cells, and ILCs. This limited the yield of the CONNECTOME analysis. Larger numbers of patients, and confirmatory cohorts, and longitudinal analyses will further leverage the potential of this methodology for the study of asthma. Third, the sequencing depth (∼50k reads/cell) does not allow the capture of all transcribed genes in the cells, especially low abundance transcripts. On average, we captured 2,055 genes per cell (range: 334 to 5,998 genes/cell). Recognizing there is a balance between reads per cell, the number of cells captured, the cost, the input of 10,000 cells to generate ∼2,000 transcriptomes provided the optimal balance for large-scale feasibility. Ultimately, the results show that with a relatively low sample number we were able to identify significant differences in asthma in a cost-effective manner.

In conclusion, single cell analysis of the sputum in asthma is a ground-breaking technique to non-invasively study the pathogenesis and heterogeneity of asthma, enabling the detailed analysis of cell populations, communication networks, and mediators that has transformative potential for understanding asthma, its heterogeneity and informing the personalized care of asthma.

## Contributors

G.L.C. conceived of and designed the study. Q.L., G.W., N.G., J.G., L.C. collected the samples and performed the experiments. X.Y., T.A., S.L., S.H., J.G., and J.S. analyzed the data, N.K., J.G., K.T., X.Y. and G.L.C interpreted the results and wrote the manuscript.

## Declaration of interests

We declare that we have no conflicts of interests.

## Acknowledgement

The authors thank all subjects who participated in this study.

## Funding

NIH NHLBI grants R01HL153604, R01HL161362, K25HL133599, NIAID grant UM1AI114271, and NLM grants R01LM014087, R21LM012884.

## Notes

### Competing Interest Statement

The authors have declared no competing interest.

