## Supplementary Materials for "Single-cell RNA Sequencing Analysis of Sputum Cell Transcriptomes Reveals Pathways and Communication Networks That Contribute to the Pathogenesis of Asthma"

### Sample Preparation and Processing

Supernatant was removed from the sputum samples. The remaining cell pallet was processed for cell suspension and mixed with diluted DTT solution to disperse the cells and permit reliable measurement of total and differential cell counts, supernatant indices, single cell RNA Sequencing (scRNA-seq) and both cell surface and intracellular markers. The cells were further processed by following single cell RNA sequencing protocol provided by 10x Genomics.

### Computational analysis

#### Data preprocessing

The raw sequencing reads were processed using the Cell Ranger pipeline (V3.1) from 10X Genomics to remove background Gel Beads in Emulsion (GEMs), demultiplex the data from different cells, map reads back to human genome, collapse aligned reads based on the unique molecular identifier (UMI) and count the number of UMIs (nUMI) for each gene. **Table S1** described the summary of sequencing data and mapping results. The nUMI counts of cells from different samples were merged into one matrix and input into the R package Seurat (V3.0)[1] for further data preprocessing including normalization, integration analysis to remove subject effect, clustering cells using Louvain algorithm, identifying cell cluster marker genes and data visualization using uniform manifold approximation and projection (UMAP). Data visualization with and without adjustment for subject effect using integration analysis in Seurat was shown in **Figure S1** as integrated data and original data, respectively.

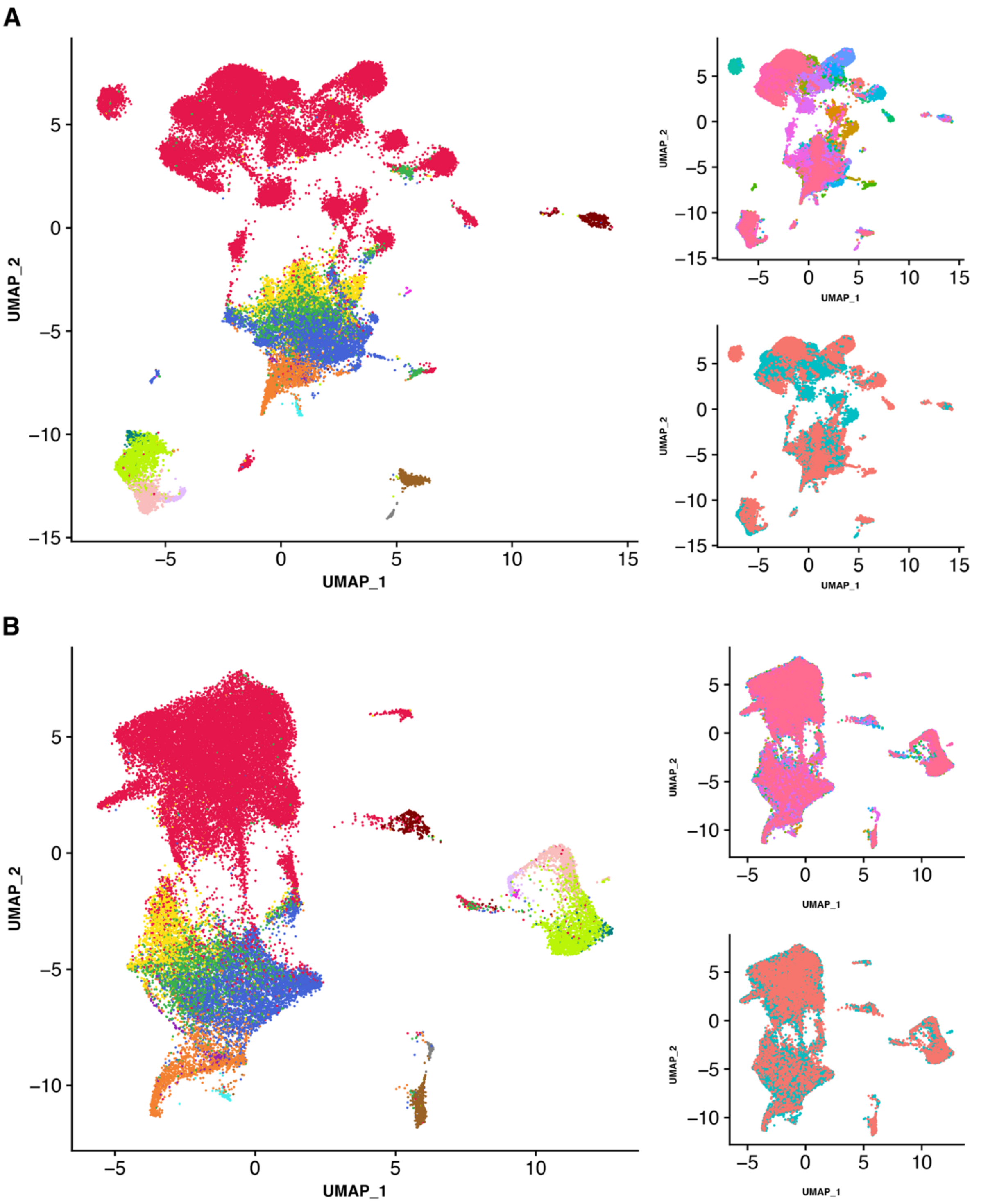


Figure **S1**: Data visualization and marker genes for the identified coarse cell populations. A) UMAPs of the original data without data integration, which were labelled using cell population (left), subjects (right top) and disease status (right bottom). B) UMAPs after the data was adjusted for subject effect, which were labelled using cell types (left), subjects (right top) and disease status (right bottom).

Difference between DMSO and fresh protocols

Among all 24 subjects, samples from 13 subjects were preserved using DMSO before being processed using the 10x protocol. For 8 of these 13 subjects, the fresh samples were also directly processed using 10x protocol without DMSO preservation (fresh). We compared DMSO preserved samples to their corresponding fresh samples from the 8 subjects to examine the difference between data of freshly processed and DMSO preserved cells from the same sample. Comparison of the distance on the UMAP between fresh and DSMO samples of the same sputum (average distance=2.51) to that between fresh samples from different subjects (average distance=5.05) demonstrated that the DMSO preserved cells are similar to the freshly processed cells from the same sputum (Wilcoxon Rank Sum test p value=0.001) than the combination of batch effect and protocol difference (**Figure S2A**). This suggests that DSMO samples are not significantly different from their corresponding fresh samples, which is consistent with previous studies[2]. Therefore, we merged the cells from DMSO and fresh protocols for the same sample together for downstream analysis (**Figure 2B**).

Dominant subject effect

Based on the visualization of the original data without integration analysis that removes subject effects, dominant subject effect existed in the data, which separated cells from different subjects from each other but made cells from the same subject cluster well together. This effect was especially high in alveolar macrophage (**Figure S1A**). Therefore, we removed the subject effects using the integration analysis by Seurat for the analysis to identify cell identity (**Figure S1B**). Note that differential expression analysis to find disease associated genes and cell type marker genes used the original data without data integration.

Missing eosinophils and neutrophils

Based on immune cell differentials measured by CytoSpin, a low percentage of Eosinophils and a high percentage of Neutrophils existed in the sputum samples, which were not captured in the 10X data. Adjustment in the 10X Cell Ranger pipeline to remove background GEMs did not change this. This was consistent with findings from previous scRNA-seq studies using Chromium V2 chips. Recent studies using 10X Genomics Chromium V3 chips were able to capture neutrophils in sputum from patients with cystic fibrosis[3] suggesting that this failure is chip specific. However, our 10x data was able to identify rarer cell types and cell types other than immune cells, including mast cells, B cells and airway epithelial cells. It also captured alveolar macrophage whose collection usually requires invasive procedures on patients such as bronchoscopy.

Cell identity analysis

The integrated analysis implemented in Seurat was conducted to remove subject effects and cluster the cells into 28 clusters (**Figure S2B**). Marker genes of each cluster were identified using the FindAllMarker function in Seurat, which compared the expression of each cluster to all other cells to find genes with significant increase. These marker genes were compared to those in previous publications[3-5] to assign coarse cell identity to each cell cluster. Doublets were defined if the corresponding markers include canonical marker genes for more than one cell types. The identified doublets were excluded from downstream analysis.

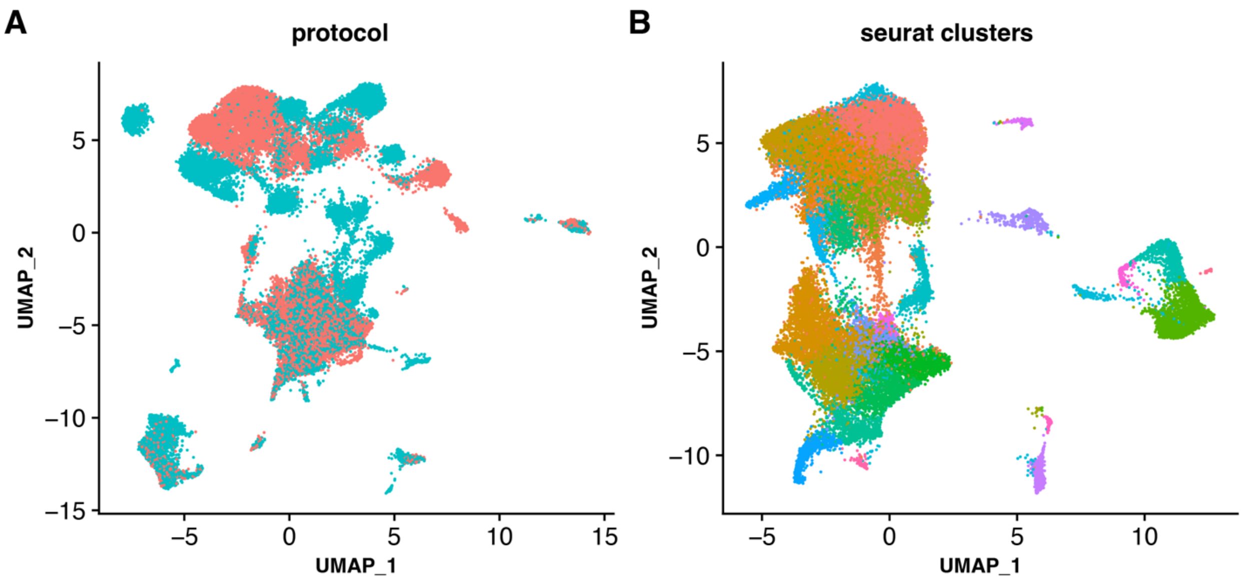


**Figure S2.** UMAPs of the A) original data labelled by protocol (DSMO or FRESH) and B) integrated data with subject effects removed labelled by the identified clusters of cells.

For coarse cell types, including T lymphocytes, B lymphocytes, macrophages, dendritic cells and airway epithelial cells, cell re-clustering was performed to identify refined cell subtypes using the same approach as that used for coarse cell type identification. The identified refined cell subtypes and their marker genes within each coarse cell type can be found in Figure **S3**. Specifically, within airway epithelial cells, cell re-clustering identified 11 sub-clusters of cells, among which 2 clusters containing 83 cells were found to have high expression of the SPRR genes (SPRR2F, SPRR2C, SPRR2B) and TGM3. These 2 clusters were squamous cells and excluded from further downstream analysis to focus our analysis on airway cells.

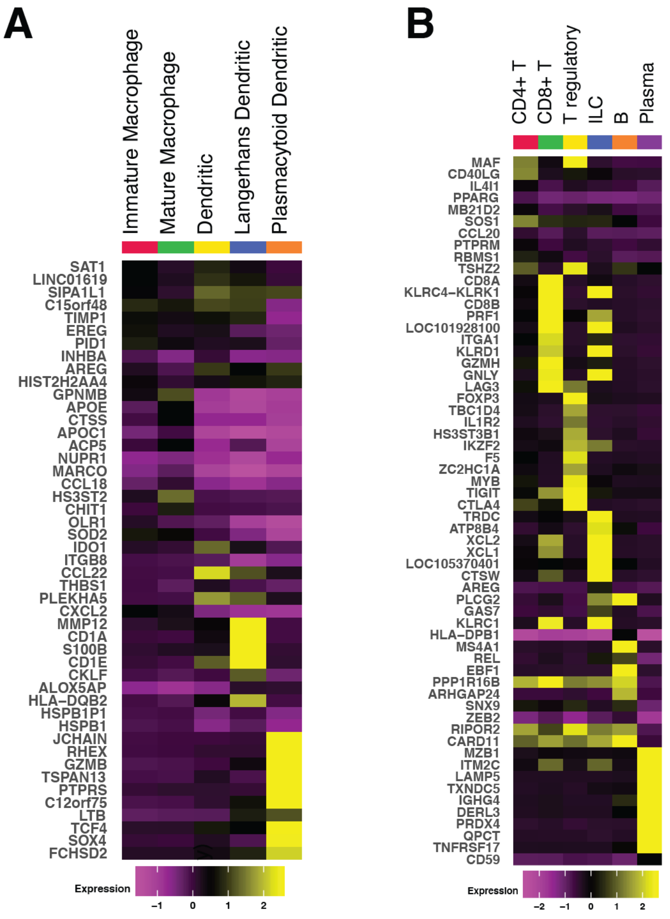


Figure **S3**. Heatmap of marker genes of refined cell subtypes within the (A) myeloid cells and (B) lymphoid cells.

<u>Cell type proportion difference between asthma and control</u>

We also compared the cell type proportion between asthma patients and controls. The most significant cell types were alveolar macrophages (p=0.04) and airway epithelial cells (p=0.06) although the difference for airway epithelial cells was marginally significant. Asthma patients tended to have higher proportion of airway epithelial cells and lower proportion of alveolar macrophage (Figure **S4**). The higher percentage of epithelial cells was potentially due to the airway epithelial shredding in asthma patients [6,7].

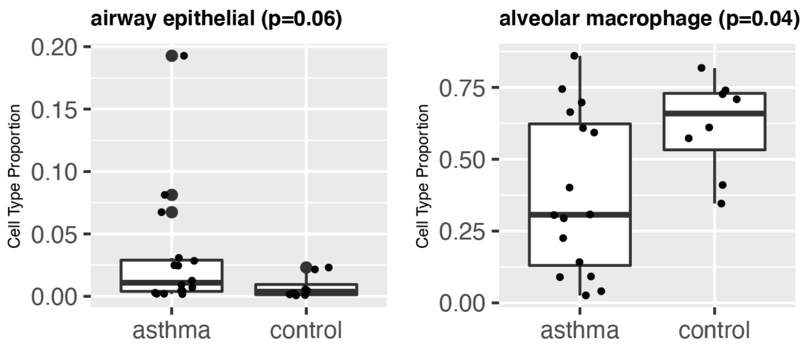


Figure **S4**: Boxplots of the cell type proportion of airway epithelial cells (left panel) and alveolar macrophages (right panel) in asthma and control. The significance of the difference (p) is shown on the top of each panel.

<u>Connectome analysis</u>

We infer the cell cell communication networks using Connectome and Fantom5 database. The inferred networks (Figure **S5**) showed intense communications between airway epithelial cells and other cell types in both control and asthma despite of the small number of airway epithelial cells, especially in controls. We calculated the incoming and outgoing centrality of each cell type in both asthma patients and controls using the inferred networks and assess the significance of difference (Figure **S6**).

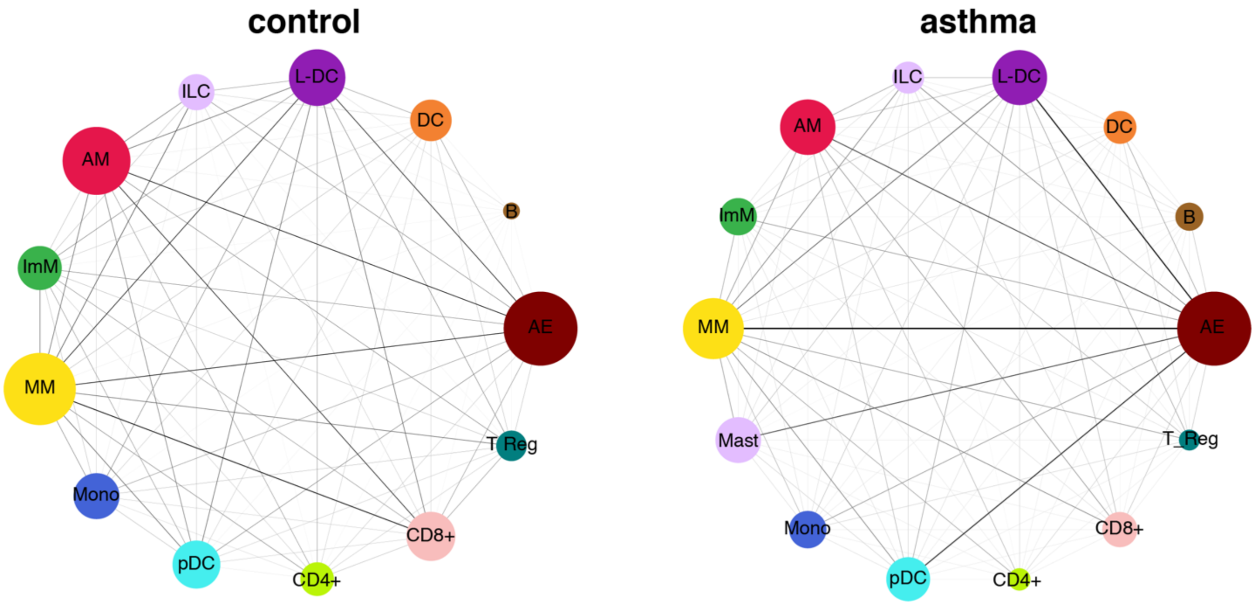


Figure **S5**: Visualization of the inferred cell-cell interaction networks in control and asthma.

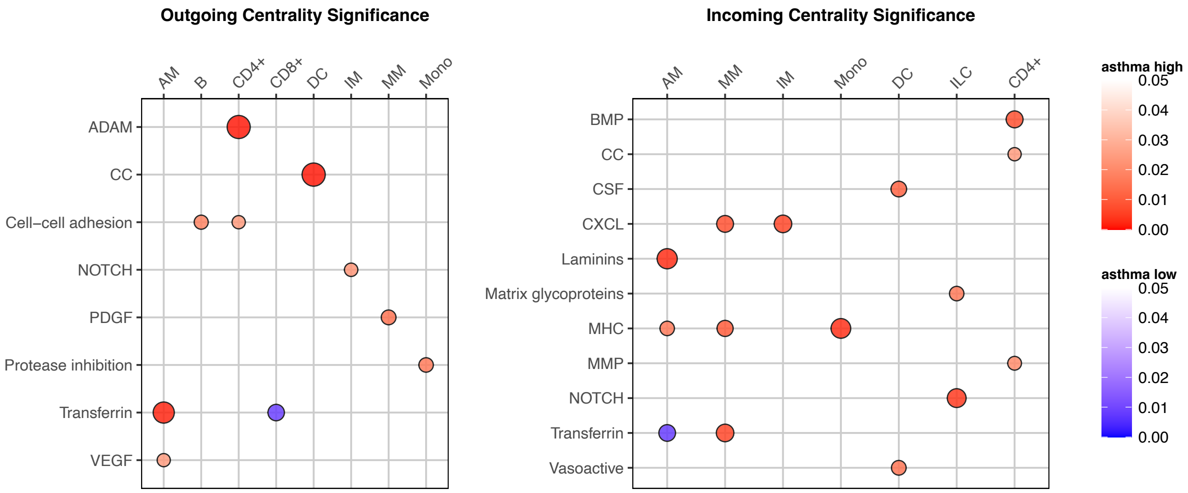

Figure **S6**: Bubble plots demonstrating all significantly (p value<0.05) different outgoing and incoming centrality between asthma and control. Red represents higher centrality in asthma and blue represents lower centrality in asthma.

Capturing Eosinophils using Chromium Single Cell Expression Flex

We collected 2 whole blood, 1 sputum sample, and 1 mixed blood and sputum sample from 2 individuals. These samples were processed using the Flex Gene Expression assay from 10x Genomics to reserve fragile cell types like eosinophils. White blood cells were isolated using buffy coat from STEMCELL Technologies INC from Cambridge, MA. The isolated blood cells were processed using the same protocol as what was used to process cells from the sputum samples.

The sequencing data was processed using the same pipeline that was used for the Chromium Single Cell 3' Reagent Kit data. In total, 96,317 cells were captured which were clustered into 25 clusters (Figure **S7A**). Azimuth was used to identify the type of each cluster using reference data from both human whole lung data and our sputum scRNA-seq data measured by the 3' kit. One of the clusters, cluster 18 (Figure **S7A**), was partially annotated as mast cells, T cells and B cells by Azimuth using a previous scRNA-seq data from human lung samples as reference data. Marker genes of this cluster from our data include HDC, GATA2, IL3RA, FCER1A, CLC and CCR3 (Figure **S7B**), which are different from the top marker genes of mast, T or B cells in the reference data. In addition, the top marker genes of mast cells, T and B cells in the reference data do not have significantly higher expression in cluster 18. Taken together, we believe that this cluster is the eosinophil population instead of mast, T or B cells.

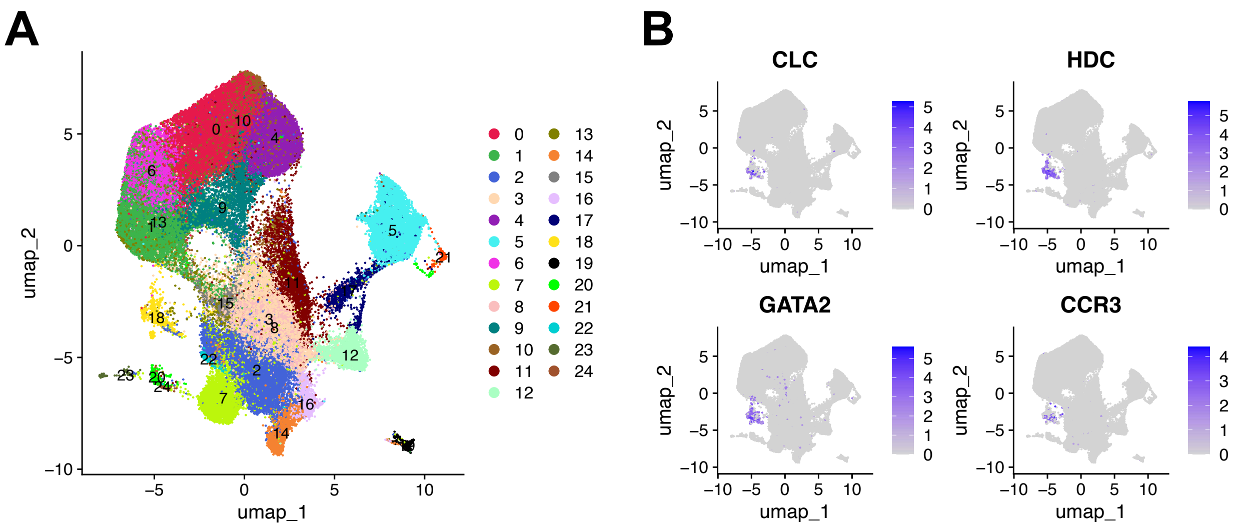


Figure **S7.** UMAP of cells captured by the 10x Genomic Flex kit in the 2 whole blood, 1 sputum and 1 mixed blood and sputum samples.
